# Human escape in freely moving virtual reality follows a structured movement pattern shaped by threat and context

**DOI:** 10.1101/2025.07.27.662958

**Authors:** Yonatan Hutabarat, Juliana K. Sporrer, Jack Brookes, Sajjad Zabbah, Lukas Kornemann, Paolo Domenici, Dominik R. Bach

## Abstract

Evading danger is critical to survival. In non-human animals, escape strategies are shaped by neural, biomechanical, and ecological constraints, resulting in species-specific patterns. In humans, ethical and practical constraints have until recently hindered investigation of escape movements, such that its organising principles are commonly extrapolated from other, mostly quadruped, species. Here, we use freely moving virtual reality (fm-VR) in a large physical space to present biologically relevant threats. We discover that human escape behaviour is organized within a constrained action space shaped by threat and context, revealing a small set of stereotyped kinematic patterns that are not predicted from those reported in other mammals. Pattern selection is shaped by environment and individual preference, and is not predicted by behaviour in safe conditions. The dominant kinematic pattern includes head orientation toward the threat, body rotation beyond this angle until facing away, and escape initiation with ipsilateral foot. Alternative variants include turning away from the threat, backward movement, and misdirected flight. Certain alternative escape patterns and kinematic features reduce success, whereas specific preparatory adjustments enhance it. Our results provide a foundation for probing the neural mechanisms of human escape, and enable investigation of potential disruptions in clinical conditions.

## Introduction

Survival across the animal kingdom depends critically on the ability to detect and evade life-threatening danger, such as predator attacks^1–3^. Animals rapidly deploy complex and often situationally flexible defensive behaviors^1,2,4,5^, which are shaped by environmental demands including features of the opponent^6–8^, biomechanical characteristics of the agent^9,10^, and the capabilities of its neural system^11^. While broad categories, such as the fight-flight dichotomy, are conserved across species, the specific expression of escape behavior is highly diverse. As a result, even closely related species can exhibit markedly different motor strategies when escaping^12,13^. Decades of field observations and laboratory experiments have mapped the structure and variability of escape responses in many non-human animals^3–5,11,14^. Human behaviour is likely to differ from that of other animals both due to their unique cognitive capabilities and specific biomechanics shaped by bipedal gait. However, ethical and practical constraints have precluded empirical investigations in humans. This is a significant gap, given longstanding speculation that individual differences in the neural mechanisms governing defensive behavior contribute to psychiatric vulnerability—particularly for anxiety and affective disorders^15^. To investigate such risk factors, studies have relied on simplified, screen-based tasks, typically mimicking escape behavior in an abstract manner by button presses or joystick movements^16^, while the characteristics of actual escape movements in humans remained unknown.

Recent advances in immersive, fm-VR and motion-tracking technology now allow expanding beyond these tasks and enable the study of realistic human behavior in simulated environments^17–21^. Here, we leverage a fm-VR platform that simulates biologically relevant threats to humans, including apex predators, aggressive non-predators, and aggressive conspecifics^18,22,23^. We investigate human movement patterns under direct attack from fast-moving opponents (figure 1) in the presence of a single shelter for survival. Using a data-driven approach with a rigorous exploration-confirmation design, we find strong evidence, replicated across three experiments (E1-E3) with overall N = 95 individuals, for a dominant escape-to-shelter sequence. We then identify consistent predictors of alternative sequences, the relation of escape kinematics and escape success, and individual preferences in its implementation.

**Fig. 1.**
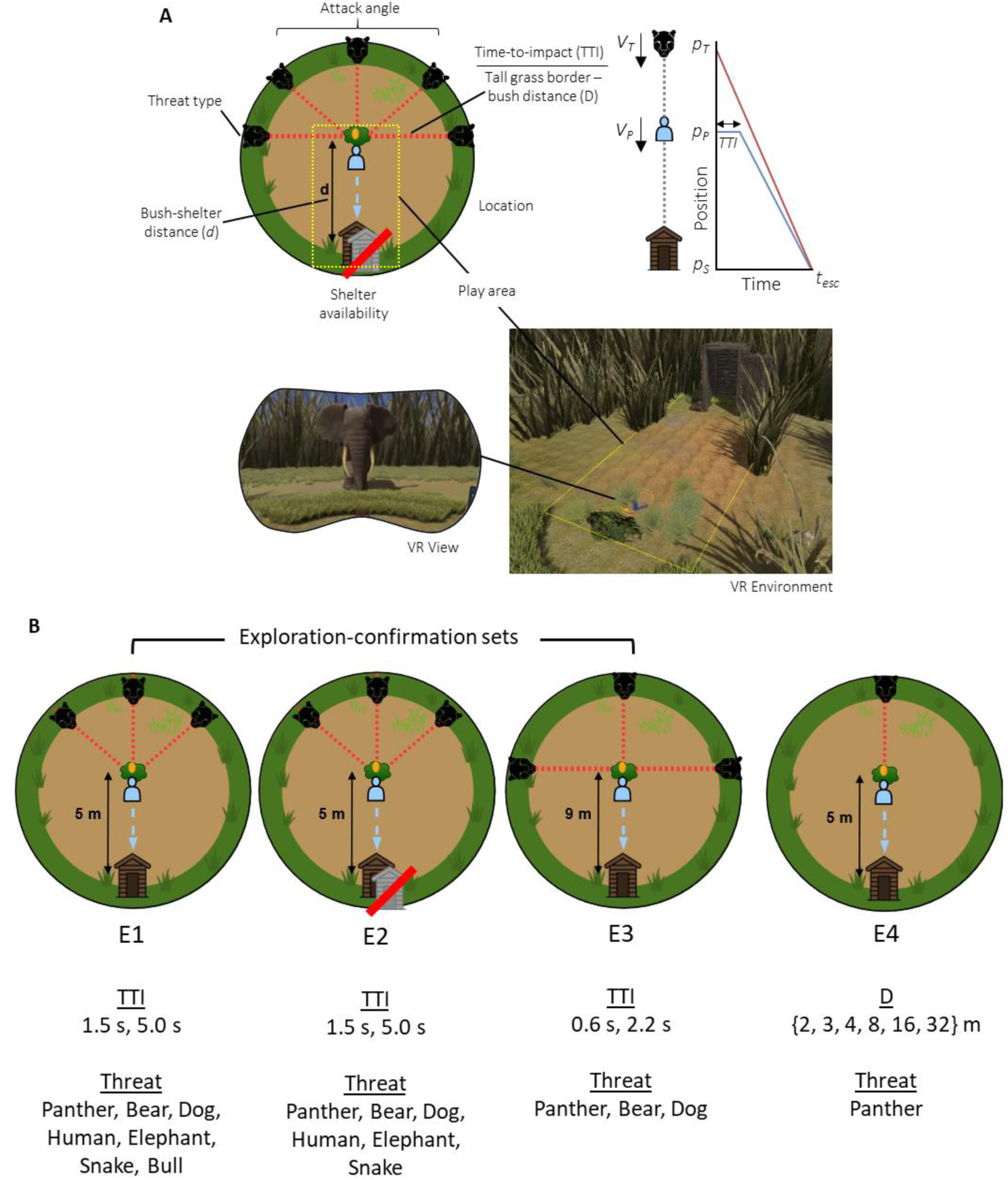
Overview of the experimental design. (A) Schematic of the experimental setup and key parameters. Four experiments (E1-E4) were conducted across two sites: University College London, UK for (E1, E2, E4; physical area of 5 x 10 m) and University of Bonn, Germany (E3; physical area of 7 x 17 m). Manipulated parameters included attack angle, time-to-impact (TTI), threat type, shelter availability, and bush-shelter distance. TTI^22^ is defined as the maximum time available to initiate escape after threat appearance at an assumed mean escape speed (*V_P_*) of 2 m/s and just about colliding with the threat at shelter entrance at t = t_esc_. Threat speeds (*V_T_*) are listed in Table S1. (B) Summary of experimental conditions across all experiment. E1 was designed for exploration, E2 for direct replication, and E3 for conceptual replication at a different site with wider attack angle and longer escape distance. E4 served to answer questions following from E1-3 analysis.

## Results

### Dominant movement sequence to fixed shelter location

First, we analyzed how participants evaded a fast-approaching opponent by moving to a shelter located 5 m behind them (515 successful escape epochs in 58 participants across E1/E2). Participants were instructed to forage as many fruits as possible in a simulated foraging task, while avoiding contact with threats. No further instruction was given, and there was no explicit incentive to forage. Participants experienced a safe shelter made of raw wood with self-closing door in a preceding tutorial, which was present in all epochs. Each epoch began with the participant moving towards a low fruit bush on a grass clearing bordered by visually occlusive tall grass, in order to pick fruit from this bush. While collecting fruit, without warning, an opponent would emerge from the tall grass, randomly at one of three angles relative to the participant-shelter axis: −45° (left), 0° (straight ahead) or 45° (right) (Figure 2A). The opponent would immediately initiate pursuit, and attack in a biologically realistic manner once it reached the participant. Contact of the threat with any body part led to virtual death, while entering any body part into the safe house lead to closure of the shelter door and virtual survival. In both cases, the epoch ended immediately, with a short display fade-out time. Because different opponent species (e.g., leopard vs. dog) move at very different speeds (Table S1), distance between the emerging threat and the fruit bush was calibrated such that maximum time to leave the fruit bush and start escape (time to impact, TTI) was 1.5 s or 5.0 s based on a realistic mean escape speed of 2 m/s. Within each experimental block, the order of epoch types was independently randomized for each participant, thus effectively minimizing potential trial order effects (about which we had no hypotheses) on group-level results. Exploratory analysis showed no differences between threats or order effects unless explicitly stated.

**Fig. 2.**
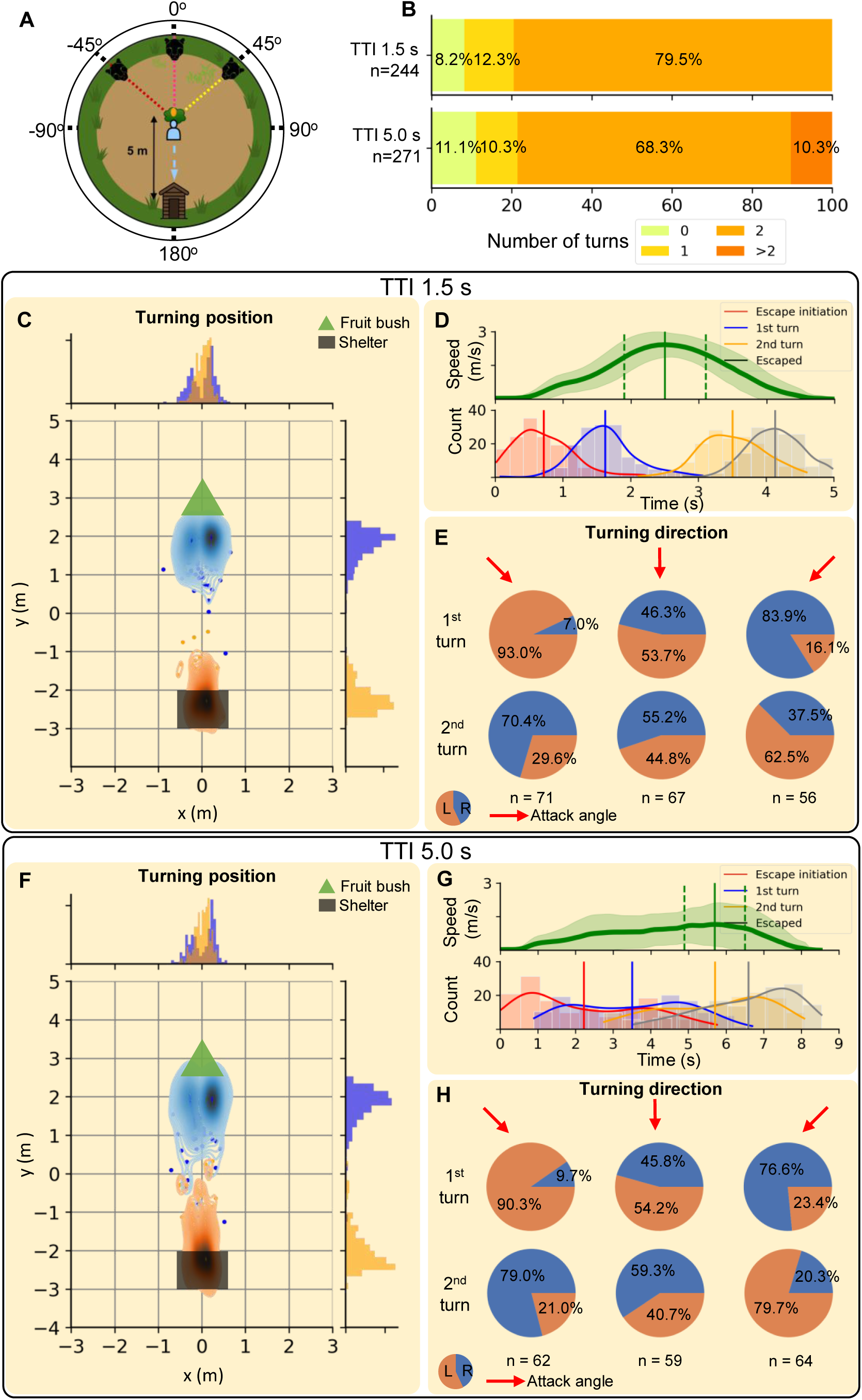
Temporal and spatial characteristics of escape to shelter (E1/E2). (A) Schematic view of the scenario setup. Attack angle is from left (−45°), front (0°) or right (45°). (B) Number of body turns (beyond x-axis) after threat appearance. The dominant movement sequence entails two body turns. (C, F) Distribution of the turning position (position of the waist tracker located on the back of the participant at the time point of turning beyond the x-axis), pooled across all attack angles. Blue points/density shows the first turn and orange the second turn. (D, G) Speed profile and temporal sequence of events following threat appearance at *t* = 0 s, pooled across all attack angles. The speed profile shows mean (thick green line) and ± 1 SD (shaded area), as well as mean timing of peak speed (vertical line) and its standard deviation (vertical dashed line). For the event sequence, thick vertical lines represent means, and the distribution represents a kernel density estimate. Color-coded events are: escape latency (red), first body turn (blue), second body turn (yellow), and successful entry into the safehouse (black). (E, H) Proportion of turning direction in relation to threat attack angles (red arrow at −45°, 0°, 45°) for the first turn (first row) and the second turn (second row). L indicates left body turn, R indicates right body turn, and n indicates number of trials for each condition. Panel C, D, E for TTI 1.5 s, and F, G, H for TTI 5.0 s.

On 79.5%/68.3% of epochs (for TTI 1.5 s or 5.0 s, respectively, Figure 2B), participants exhibited a distinct escape sequence (Figure 2, Supplementary video). This was characterized by movement initiation, head orientation toward the threat, body turn in the same direction and then beyond 90° relative to fruit-shelter axis, forward locomotion towards the shelter with a (mean±SD) peak acceleration of 5.08±1.18/4.35±1.32 m/s^2^ and peak speed of 2.85±0.39/2.40±0.54 m/s, respectively, followed by a final body turn towards the threat before entering the shelter backwards (Figure 2CDFG). Location of the turning positions are shown in Figure 2CF. The distributions of head and body orientation over time (Figure S1) demonstrate that body orientation followed head orientation during the initial seconds after the threat appeared. Interestingly, escape latency had a wide distribution, including values below 200 ms (Figure 2DG).

Crucially, when an opponent attacks laterally, any escape direction can be reached by initially turning away, or toward the threat. In this situation, many animals prefer to turn away from the threat^1^. Human participants, in contrast, predominantly turned their head and then their full body toward the threat (Figure S1), then kept turning in the same direction until the body was approximately facing the shelter (Figure S1), at which point they started running forward. Thus, the first turn (beyond 90° relative to bush-shelter axis) was in the direction of the threat in more than 84.2% of epochs (p < .0001; Figure 2EH, full statistical results divided by E1-E3 in Table S2). Notably, this turning direction preference showed some degree of lateralization: it was more pronounced when the opponent attacked from the left (−45°) than from the right (45°; p < .0001; Table S2).

Next, we were interested to test if this escape pattern was affected by environmental parameters. In E3, opponents attacked from an angle of −90°, 0°, or 90°; TTI was 0.6 s or 2.2 s; and the shelter was 9 m away. Again, we analyzed epochs with successful escape from a fast-approaching opponent (Table S1) that emerged and attacked without warning (191 successful escape epochs in 36 participants). On 70.0%/83.2% (TTI 0.6 s or 2.2 s, respectively) participants exhibited the previously found escape sequence (Figure S2), with the first turn predominantly toward the threat (p < .0001), especially when the attack was from the left (p < .0001). Mean peak acceleration and mean peak speed were 5.66±1.27/5.31±1.16 m/s^2^ and 4.00±0.43/3.70±0.52 m/s, respectively.

### Deviations from dominant movement sequence during successful escape to shelter

Next, we focused on deviations from the dominant movement sequence in epochs with successful escape from a fast-approaching opponent. In 9.7% of these epochs in E1/E2, participants did not turn during successful escape, and instead moved backwards to the shelter (Figure 2B). This never happened in E3, where the shelter distance was 9 m instead of 5 m (Figure S2).

In 11.3% of epochs in E1/E2 and 23.0% in E3, only one body turn was observed, which took place within 2 m from the foraging area in 94.8% (E1/E2) or 93.2 % (E3) of instances and corresponded to the predominant movement sequence without final turn at the shelter. Finally, in 5.4% of epochs in E1/E2, participants turned more than twice. This occurred only at a TTI of 5.0 s in E1/E2 and never happened in E3, where TTI was 2.2 s or smaller.

### Threat type and individual characteristics influence escape strategy

Interestingly, in E1/E2, the type of opponent influenced whether participants would move backwards or not. For this analysis, we combined successful with unsuccessful escape attempts to avoid bias. We observed more backwards escape attempts for dog, snake and human opponents (overall 23.2 %) than for bear and elephant (overall 8.8 %, p < .05; Table S2, Figure S3). At the individual level, most participants followed the dominant escape sequence most of the time, however some participants displayed a strong preference for backward escape (Figure 3A). Furthermore, while many participants followed the pattern of a predominant turn towards the threat, a small number consistently deviated from this pattern and showed a strong preference for turning predominantly either to the left or to the right, consistently for both attack directions. No participant consistently turned away from the threat for both attack directions (Figure 3B).

**Fig. 3.**
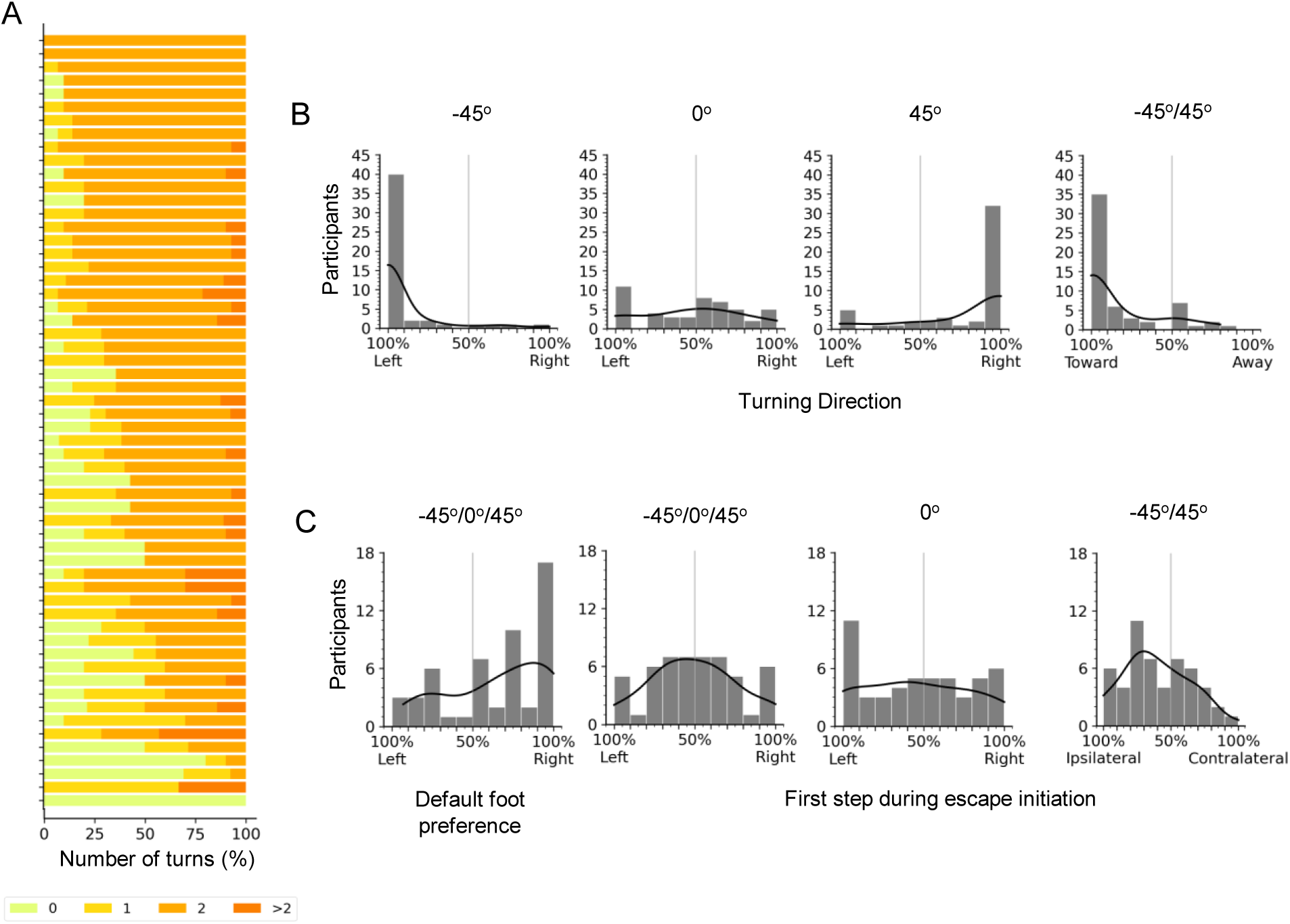
Turning direction and foot preference (E1/E2). (A) Individual turning counts for all participants and all attack trials, sorted by proportion of dominant movement sequence (2 turns) for all TTIs. (B) Turning preferences for individual participants, for all attack trials and TTIs, separated by threat attack angle. No participant consistently turns away from the threat (panel - 45°/45°), showing that the deviations from dominant turning direction seen for individual attack angles are due to a direction preference. (C) Default foot preference for individual participants for all attack trials, assessed during first step initiation without threat; and foot preference for individual participants during escape initiation.

### Determinants of unsuccessful escape

We recorded 155 (88/67 in E1/E2) epochs in which participants were caught by the threat. The most pronounced difference between unsuccessful and successful escape was the speed profile (Figure 4A, Table S3) with slower mean speed (p < .0001 for both TTI) and peak speed (p < .001 for both TTI), lower mean acceleration magnitude (p < .0001 for both TTI), lower peak acceleration magnitude at short TTI (p < .05), and later peak speed and later peak acceleration at long TTI (p < .01). Furthermore, unsuccessful escape was initiated later than successful escape (p < .001 for both TTI, Figure 4B, Table S3), although in most instances, participants could still have escaped successfully at an escape speed within the range observed. Finally, participants were around four times more likely not to turn at all in unsuccessful escape attempts (Figure 4C). To exclude that this was simply due to getting caught before they could turn, we analyzed the time when they were caught in these instances (4.01 ± 0.81 s/7.23 ± 0.85 s), which was much later (p < .0001, Table S2) than the average first turn time in successful escape (1.64 ± 0.46 s/3.31 ± 1.51 s). Thus, it appears that participants intentionally moved backwards without turning, and that this contributed to their unsuccessful escape. Finally, as shown in Figure 4D, there were many instances of misdirected flight (12.5%/8.9% for TTI 1.5 s/ TTI 5.0 s), in which participants did not run towards the shelter, and instead towards the tall grass, where they would get caught by the opponent. This occurred throughout the experimental session (median trial = 14.5; Interquartile range (IQR) = 23). Interestingly, 40% (12/30) of all misdirected flights occurred upon encountering a snake (113 of trials).

**Fig. 4.**
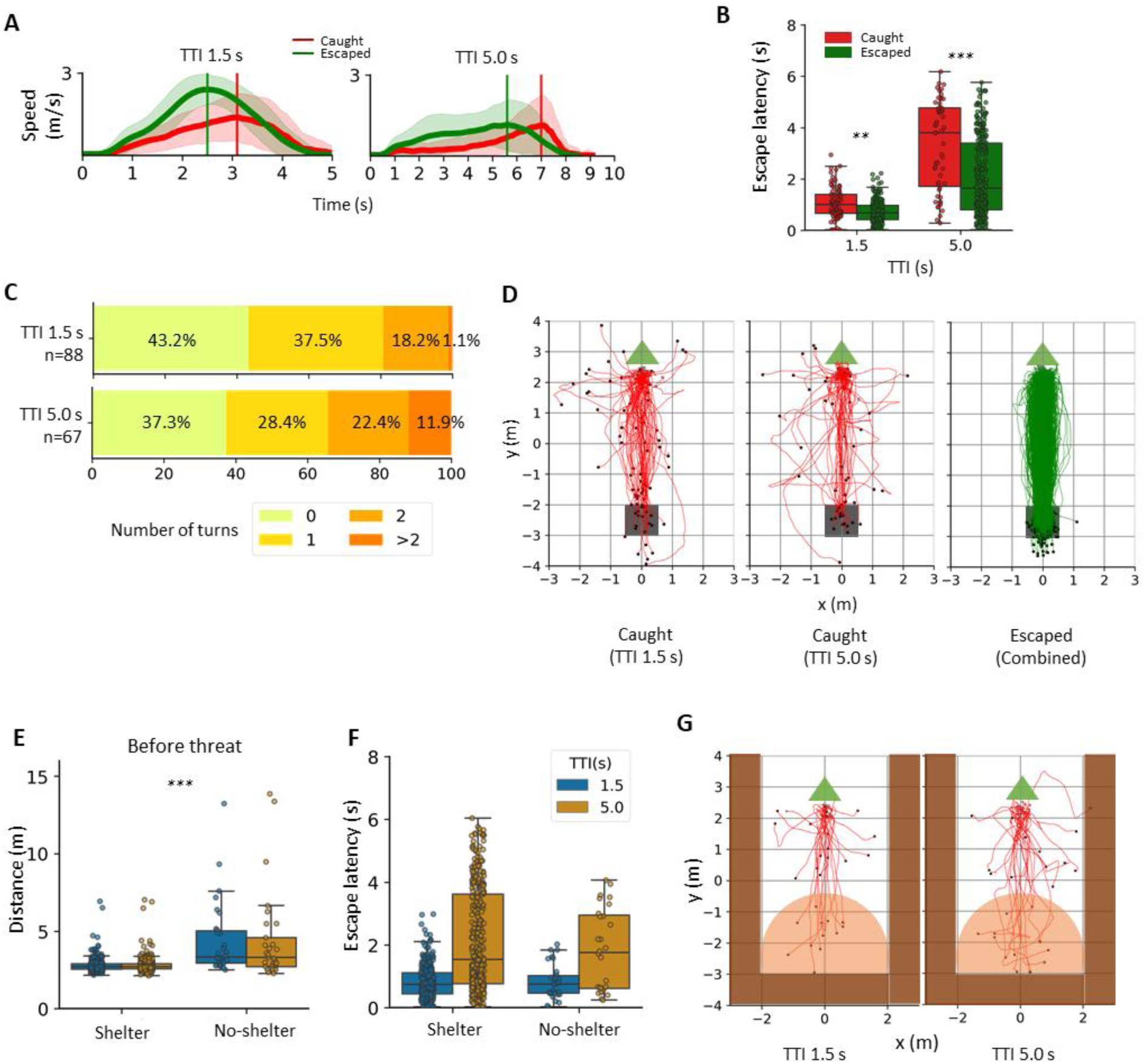
Determinants of unsuccessful escape and escape in absence of shelter. (A) Speed profile for successful and unsuccessful escape to shelter (B) Later latency for successful and unsuccessful escape to shelter. (C) Turning counts for unsuccessful escape to shelter (see figure 1 for successful escape). (D) Escape trajectories for unsuccessful escape (red), with instances of escape to non-shelter directions, compared to trajectory of successful escape (green). Black dots: end position where participants were caught/escaped. (E) Distance walked before threat appearance and (F) escape latency for epochs with known shelter location vs. no-shelter. (G) Trajectory in no-shelter epochs, showing that escape is often towards the previous location of the shelter. The ravine is depicted in brown surrounding the play area, and the gray area indicates a 2 m radius from the location of the previously available shelter. Panels ABEF show mean ± SD. *\*\* p < .001, *** p < .0001*.

### Defensive efficiency is prioritized over habitual movement patterns

Humans commonly initiate locomotion with a preferred foot. At the start of the epochs, when participants had to move towards the fruit bush in the absence of any threat, the majority of participants initiated walking with the right foot (p < .0001; E1-3, Table S4, Figure 3C). In contrast, when initiating escape, foot preference was reduced (p < .001, Table S4), including when threat attack was from a 0° angle and thus turning direction was not externally influenced (p < .01; Table S4). A mixed-effects logistic regression analysis with threat attack direction (−45°/-90° vs. 45°/90°), turning direction (left vs. right) and initial foot preference (left vs. right) as predictors showed that escape foot preference was directly determined by turning direction (p < .01; Table S5, Figure 3C), with no evidence for an additional impact of threat direction (p = .70) or default foot preference (p = .48). These findings suggest that while threat direction affects turning direction, it has no direct impact on foot preference. Taken together, these findings suggest that default motor preferences from non-threatening situations are overridden by situational demands, which mainly depend on the chosen turning direction.

### Turning direction is predominantly non-protean

Some animals use protean, i.e. unpredictable, defensive behavior, to protect against predators^24–26^. Unpredictability can also arise at the population level if individuals have stable preferences but these are randomly distributed in the population^5^. On the other hand, some species exhibit strong preferences for a specific turning direction even at the group level^1,27,28^. Here, we sought to determine whether turning direction in humans in repeated encounters with the same threat followed a protean or non-protean strategy when attacked in a non-directional (i.e. head-on) manner. Data set E4 included a block of consecutive epochs with head-on attack (0° attack angle) from the same threat (black-coloured leopard, 561 epochs in 47 participants). We tested whether individual participants had random or non-random sequences of turning directions. At the group level, we found strong evidence against sequence randomness (Wald-Wolfowitz test with Fisher’s method^29,30^: χ^2^= 275.03, df=80, p < .0001), and 27/47 participants even turned deterministically into the same direction. Taken together, this suggests that human turning patterns to frontal attack are predominantly non-protean at the subject level.

### Escape in absence of shelter

Next, we examined epochs without shelter, situated in a ravine (Figure S4) without plausible exit routes (57 epochs in 30 participants, E2 only), which always took place after participants were accustomed to the fixed shelter location. Before the threat appeared, participants covered more distance compared to epochs with known and fixed shelter location at the same timing (Figure 4E, Table S2). Once the threat emerged, escape initiation time was not statistically different from trials in which the shelter was available (Figure 4F). Finally, participants approached the previously available shelter location in 41%/50% epochs, even though the shelter was absent (Figure 4G), reminiscent of similar behavior in mice^31^.

### Distinct foraging posture at short threat distance predicts successful escape

E4 included instances in which a black-coloured leopard emerged from tall grass at much shorter distance (2-4 m) than in E1/E2 (minimum 8.8 m). Here, we observed a notable shift in foraging strategy before threat encounter. At close proximity to the edge of the grass area, participants altered their stance by widening their base of support (Figure 5A). Specifically, the distal foot was positioned away from the proximal foot (adjacent to the bush), thus lowering the center of gravity as well as head position which creates a more consistent visual pathway that spans both the bush and the forward tall grass, providing a broader horizontal scanning ability without frequent head reorientation in preparation for escape.

**Fig. 5.**
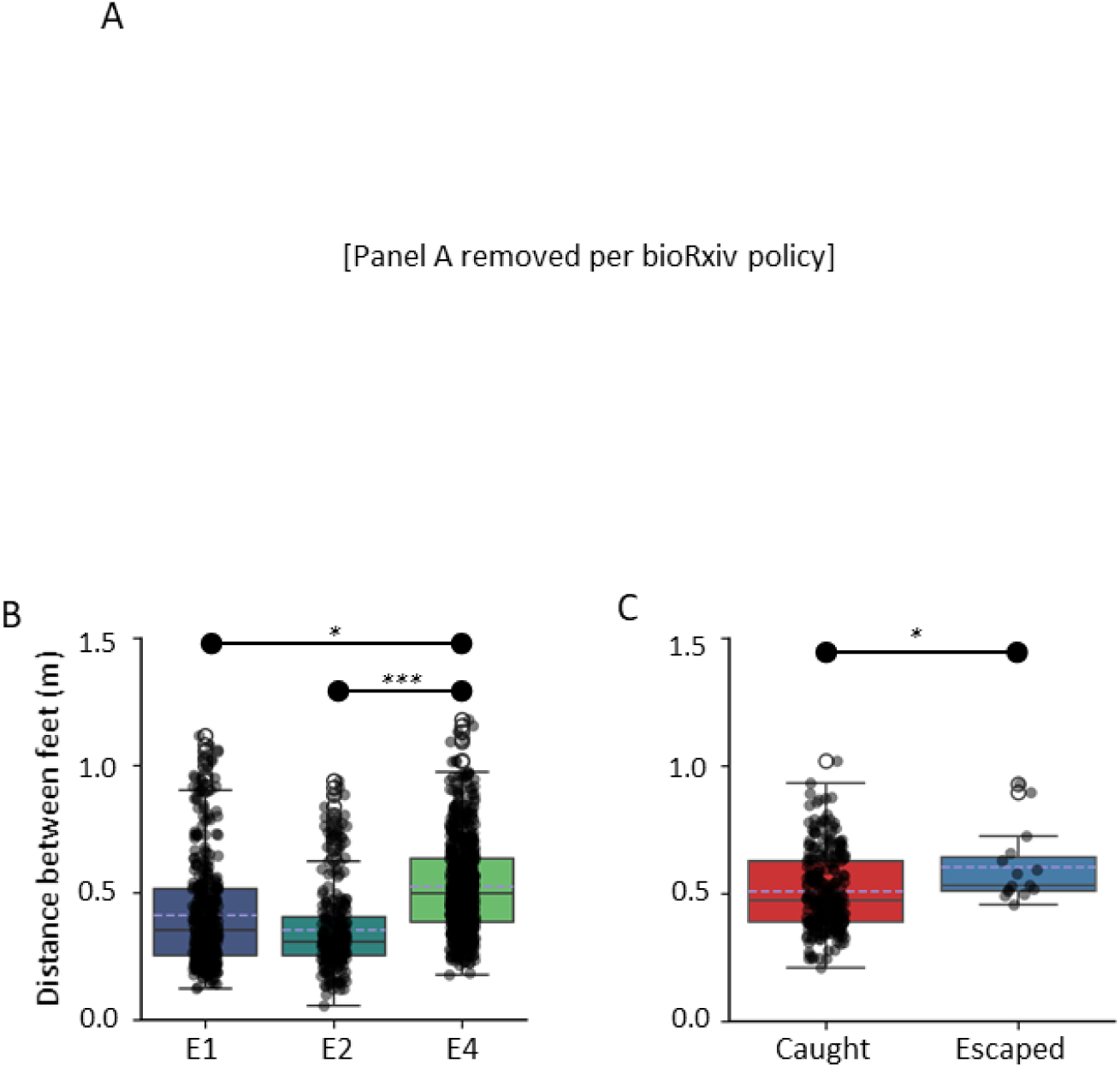
Foraging strategy in E4. (A) Multi-camera view of a participant foraging on the bushes, where wider stance is favored for escape preparation. (B) Comparison of foraging posture from E1, E2, and E4 as indexed by Euclidian distance between participants’ feet sampled from the first second of first fruit collection. (C) Comparison between successful and unsuccessful escape at close grass-bush distances (2-4 m) in E4. Dashed purple line indicates the mean of each group. Panels BC show mean ± SD*. *p < .01, *** p < .0001*

When the threat appeared, the foot closer to the bush responded rapidly, generating an impulse to move away from the bush, while the distal leg maintained support. This dynamic response facilitated a swift relocation of the center of gravity toward a safer direction. We found that this strategy differed significantly from what we observed in E1/E2, where distance between participants’ feet was smaller (p < .01; Figure 5B, Table S6). Furthermore, at grass–bush distances of 2-4 m, distance between feet predicted success of escape (p < .01; Figure 5C, Table S6).

Interestingly, the postural adjustment during fruit collection (Figure 5A), where left foot was positioned furthest from the bush, was observed in the majority of trials (72.1%, n=725), showing a distinct population-level lateralization on foraging posture.

## Discussion

A body of field and laboratory evidence indicates that many animals employ species-specific behavioral patterns to evade predators and aggressive conspecifics^5,7,8,11^. Here, we use fm-VR with realistically simulated biological threats, to investigate movement sequence in humans. Our findings provide four key insights.

First, humans exhibit a characteristic escape-to-shelter movement sequence most of the time. The sequence included an initial head turn towards the threat, followed by a body turn in the same direction until facing shelter. At this point, participants started running toward the shelter using their ipsilateral foot first. We also observed a final turn at the shelter entrance marking escape termination^4^, where participants transition from forward locomotion a monitoring state. Phenomenologically, this resembles flight to hide behavior in prey animals, which immediately reorient at the entrance to monitor and defend against a predator^32^. Notable deviations from this pattern included backwards escape, which occurred at varying rates in different participants, and was more often associated with unsuccessful escape attempts. This underscores the adaptive function of the dominant movement pattern. It is worth noting that the types of escape observed here are likely driven by short-latency responses and are geometry-dependent. Additional escape strategies may emerge in longer latency scenarios that allow more planning and with different spatial relationship between threat-participant-shelter. Conversely, other plausible deviations from characteristic escape patterns were not observed, such as preferentially turning away from the threat, a behavior observed in some animals^5^, or orienting towards shelter without regard for the threat position as observed in mice^34,35^. We suggest that human turning preference may reflect certain characteristics of the sensory system, such as the eyes located in front rather than at the sides of the head as in some other animals. At the individual level, no unpredictability in turning direction was observed; instead, we observed strong individual preferences when attack direction was head-on, and a few individuals showed turning preferences even when attack was lateralised (Figure 3B). At the group level, there was a slight preference for left turns. It is worth noting that many vertebrates with laterally placed eyes exhibit a consistent lateralization of preferring to monitor potential threat with the left eye^36,37^, and there is a suggestion that such lateralisation might confer an adaptive advantage in dual-task settings such as foraging under threat^38,39^. Given the different eye position in humans, future research will address whether the turning preferences observed here correspond to laterality of threat monitoring.

Second, we found that typical movement patterns in non-threatening situations were altered in the presence of a threat. Specifically, person-specific foot preference, as observed before the appearance of a threat, did not predict foot preference under attack. Here, foot usage was exclusively determined by the selected turning direction. This finding highlights the optimization of action execution under threat.

Third, in the absence of a shelter, participants explored the environment more than in the presence of a shelter during the pre-encounter phase, but escaped with similar timing once a threat appeared, often to the position of the previously available shelter. Similar behavior is observed in mice who locomote to previously available shelters even if they have been removed^31^.

Finally, at very short threat distances, participants increased their foot distance, which lowers the center of mass and increases the base of support, during the pre-encounter phase to one that potentially facilitates a more rapid thrust away from the threat, while also enhancing stability during the initial acceleration. This strategy provides better leverage and control over torques generated around the hip and trunk^40–42^, optimizing the push-off phase and horizontal thrusting to the direction of shelter. Moreover, we observed population-level lateralization in foraging posture such that left foot was positioned further away from the bush, suggesting that under conditions of high threat predictability (known attack direction), the motor system prioritize immediate acceleration potential over the flexibility required for omnidirectional monitoring.

Escape success depends on many factors, such as how quickly a threat can be detected, the availability of a refuge, and the trajectory of escape^1,4,9,43^. Recent research emphasizes that escape behaviors integrate sensory information, memory, and internal state of the animal to produce a flexible, context-dependent response^4^. In principle, animals will attempt to put distance between themselves and a predator as quickly as possible^11^. Field and laboratory studies show that if a refuge is available and known to the animal, escape trajectories are often directed straight toward it^5^, including in mammal species such as gerbils^33^ and mice^34^. We find that in humans, unsuccessful escape is predicted by four factors: backward movement, misdirected movement toward non-shelter location, late escape initiation, and escape kinematics. In particular, acceleration was delayed and/or attenuated on unsuccessful escape attempts. In contrast, during successful escape, latencies were not infrequently below 200 ms. Across a range of studies, simple visuomotor reaction times in healthy adults cluster around 200 ms, with only small differences across age and sex^44^. However, anticipating – and presumably, preparing – a motor response can result in reaction times even smaller than 150 ms^44,45^ in various task contexts. Interestingly, misdirected flight was not exclusively observed during the first few trials, where one might argue that participants were still learning the value of shelter, but occurred predominantly when they were being pursued by the snake.

A plethora of studies have addressed how variability in human behavior towards threat might relate to the risk for psychiatric conditions^46^. This line of thinking often emphasizes the importance of testing for differences in perception, interpretation, or prioritisation of sensory information, as well as variability in decision-making under threat^15,47^. While such aspects can be investigated in third-person view computer games^16^, the latter bypass the rich motor dynamics of real-world situations, which complicate the required decision-making. The advent of fm-VR has enabled simulating real-world situations and thus, investigating actual movement patterns^18–21,48^. Using a data-driven approach, we extend these findings with a systematic analysis of human escape patterns. This paves the way towards linking such patterns with individual preferences in other domains, and ultimately, psychiatric risk.

A fundamental consideration in interpreting the findings here is the distinction between simulated and real-world threat. Non-human animals presented with simulated threat such as visual looming stimuli^10,49,50^ potentially perceive these as a genuine risk to survival but in fact quickly learn to stop escaping over repeated encounters^51^. In contrast, human participants are likely to maintain a cognitive awareness that they are in a simulated fm-VR environment. This belief gap between the real word and its VR simulation could, in theory, alter the urgency or strategy of escape response. However, we argue that the structured motor sequence we observed likely reflects the plausible optimal escape strategy in our paradigm. First, there is no explicit incentive in our paradigm to escape, and yet participants do so with little change over many trials^22^, even showing non-instructed behaviours such as vocalisations and screams^22^. This suggests that participants perceive the fm-VR as immersive and behave naturally. Second, literature on other high-stakes simulation paradigms, such as driving and flight simulators, demonstrate generalization of phenomena observed in VR to real-world scenarios^52^. A recent study on locomotion in VR confirmed that the fundamental joint kinematics and coordination patterns remain largely preserved compared to real-world scenarios^53^. A more specific limitation is that our threat characters, while realistically animated and responsive to the human, might be perceived as less intelligent than a real opponent, which might influence game-theoretic strategies such as protean behaviour.

Escaping from a threat demands an immediate response marked by high speed and precision—what has been referred to as “critical intelligence”^43^. Investigating the cognitive control of this behaviour holds significant promise to elucidate our understanding of its basic neurobiology^47,54^, its failures in clinical conditions, and potentially to inform the design of safety features in autonomous systems^43^. Central to this endeavor is the identification of the agent’s action space—the actual movements executed in response to threats. The present study marks a pivotal first step in this critical line of inquiry.

## Acknowledgements

This work was funded by the European Research Council (ERC) under the European Union’s Horizon 2020 research and innovation programme (Grant agreement No. ERC-2018 CoG-816564 ActionContraThreat) and the iBehave Network, which is sponsored by the Ministry of Culture and Science of the State of North Rhine-Westphalia, Germany. The Hertz Chair for Artificial Intelligence and Neuroscience in the Transdisciplinary Research Area Life and Health, University of Bonn, is funded as part of the Excellence Strategy of the German federal and state governments. The Wellcome Centre for Human Neuroimaging is supported by core funding from the Wellcome (203147/Z/16/Z).

## Author contributions

Conceptualization: YH, DRB

Methodology: YH, JKS, JB, SZ, LK, DRB

Investigation: YH, JKS, JB, SZ, LK, DRB

Visualization: YH

Funding acquisition: DRB

Project administration: YH, DRB

Supervision: PD, DRB

Writing – original draft: YH, DRB

Writing – review & editing: YH, JKS, SZ, LK, PD, DRB

## Competing interests

Authors declare that they have no competing interests

## Data and materials availability

All code with summary statistics used for analysis is publicly available on Open Science Framework (OSF, https://osf.io/pmdjt/). The compiled version of E1 and E2 are publicly available on OSF (https://osf.io/2b3k7/). E3 and E4 will be made publicly available upon acceptance of the manuscript. Custom code for VR threat data integration is available on GitHub (https://github.com/bachlab/vrthreat). The VR threat toolkit used to design all experimental scenarios is available under a license agreement (to protect third-party intellectual property) on UCL’s software portal XIP (https://xip.uclb.com/product/vrthreat-toolkit-for-unity). All data is available for academic purposes under a data protection agreement, due to data privacy constraints.

## Supplementary Materials

## Materials and Methods

### Participants

We report data from 29 participants (19 females and 10 males, mean age ± SD: 25.9 ± 5.6 years) in E1, 30 in E2 (15 females and 15 males, 24.7 ± 5.3 years), 36 in E3 (17 females and 19 males, 24.5 ± 4.7 years) and 56 in E4 (41 females and 15 males, 22 ± 4 years). Escape decisions from E1/E2 were analyzed previously^22^; here we report novel analyses of the same data set. All participants gave written informed consent before starting the experiment, and received a fixed monetary compensation. Experiments complied with all relevant ethical regulations and were approved by UCL Research Ethics Committee (6649/003) for E1, E2, and E4, and by the Research Ethics Committee of the Medical Faculty, University of Bonn (240/22), for E3.

### Escape elicitation

Each experiment consisted of a sequence of short encounters (‘epochs’) with various threats, and self-paced breaks in-between. Participants were tasked with collecting as many pieces of fruit on a bush (resembling kumquats) as possible, while staying clear of physical contact with any threat. Participants were instructed that to escape to safety, they could enter a shelter whenever it was available during the epoch. At the start of each epoch, participants were positioned in a low grass clearing surrounded by tall grass in a prairie-like environment, with a single fruit bush 2.5 m, or 4.5 m (E3 only), in front of them and a safe shelter 2.5 m, or 4.5 m (E3 only), behind them. Once they started fruit collection, a threat could emerge from the tall grass after a random delay. Threat appearance was accompanied by an initial rustling sound as it broke through the grass, with the audio spatialized to reflect the location of the threat. Interspersed were epochs of a different design which are not analyzed here: epochs without threat (all experiments), epochs in which the threat diverted and did not attack, epochs with slow threats that did not require escape to shelter (E1/E2), epochs in which the threat type, direction, and speed were explicitly signalled to participants (E3), and epochs in which the bush was located behind a visual obstacle such that the threat appearance was not directly visible (E4). Epochs were presented in fully randomized order. Thus, we analyzed 391, 353, 204, and 971 epochs for E1-4, respectively. For successful escape analysis, we excluded epochs where participant moved away from the fruit bush before threat appears (6 epochs in E1 and 4 epochs in E3). One participant in E2 was excluded due to faulty waist tracker (11 epochs in E2). Because of the low number virtual deaths in E3 (9 epochs), E3 data were not analyzed for the comparison of successful and unsuccessful epochs. For foot tracker data, after excluding epochs with missing or incomplete data, much fewer epochs remained for E2/3 for confirmation analysis such that we only used E1 data for foot tracker analyses.

Each threat moved with its specific, constant, and biologically plausible speed. The threats are listed in Table S1.

Threat distances were determined such that at the specific threat speed and an assumed participant escape speed of 2 m/s, the maximum time to initiate escape (time-to-impact, TTI) was fixed at 1.5 s or 5.0 s E1/E2, and 0.6 s/2.2 s in E3. The details on TTI calculation can be found in our previously published study^22^. In E4, higher participant escape speeds were required as the threat appeared at a distance of 2, 3, 4, 8, 16, and 32 m with a speed of 3.7 m/s. Threats appeared at an angle (relative to bush-shelter axis) of −45°/0°/45° (E1/E2), −90°/0°/90° (E3) or 0° (E4). Overview of all the experimental designs can be seen in figure 1.

### Movement tracking

Participants were equipped with a VR headset (HTC Vive Pro Eye HMD), two corresponding hand controllers, and three sensors on the waist and each foot (HTC Vive Tracker 2.0). Each of the six sensors yielded position and rotational information in 3D space on a frame-by-frame basis, locked to the frame rate of the VR display with a hardware-rated maximum of 90 Hz that occasionally dropped to lower rate due to GPU constraints. The HTC Vive tracking system has been extensively validated for kinematic research, demonstrating high precision and a very low average latency^55,56^. The waist sensor was positioned on the back of the participant and thus slightly behind the center of gravity, as apparent in figure 1 (i.e. when the player turns left while standing on the midline, the turning position of the waist tracker will be recorded somewhat to the right of the midline).

### Data Preprocessing

We downsampled and linearly interpolated movement data to a uniform 50 Hz rate. After performing any derivative calculations using numerical differentiation, we applied a fourth-order zero-lag Butterworth filter with a cut-off frequency of 6 Hz for smoothing. We excluded trials in which head or waist tracker data were unavailable or incomplete due to hardware failure. Additionally, we excluded trials with sudden jumps in tracker position to implausible locations in 3D space.

In the following, we define all the dependent variables reported in this paper, along with their corresponding mathematical descriptions.

### Escape latency

Escape latency denotes the interval between threat appearance and the initiation of an escape response. This metric was derived through a two-step process. First, we calculated the baseline effort, which corresponds to the average motion intensity during the initial locomotion phase (from the trial start to the foraging area). Motion intensity, I(t), for each tracker was defined as the Euclidean norm of the acceleration vector:

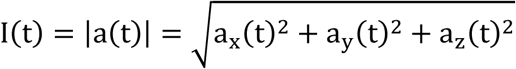

The baseline effort (*E_b_*) is then computed as:

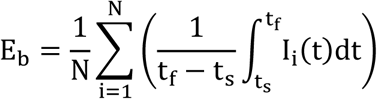

where *N* is the number of trackers, *t_s_* is the trial start time, and *t_f_* is the time of the start of foraging. Second, following the threat appearance, we computed a moving average of the effort over a 300 ms window. The escape latency was identified as the first instance when this rolling mean exceeded the baseline effort. This approach allows us to detect a sustained increase in motion intensity compared to the baseline, which we interpret as the initiation of an escape in response to the perceived threat.

### First step detection

The initiation of the escape response was characterized by detecting the first step taken after threat appearance. This process involved analyzing the acceleration data from both foot trackers. We computed the Euclidean norm (L2 norm) of the acceleration vector for each foot tracker, defined as 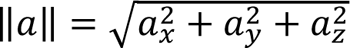 where *a_x_*, *a_y_*, and *a_z_* represent the acceleration components in three-dimensional space. A step was identified when the acceleration magnitude exceeded a predefined threshold of 3 m/s^2^. Our threshold was set above the typical acceleration range of steady-state walking (0.5 m/s^2^ - 2 m/s^2^) reported in reference ^57^ to specifically capture the more pronounced accelerations associated with step initiation, particularly in the context of an escape response. By determining which foot tracker first exceeded the threshold, we identified the initiating foot for the escape response. To assess potential lateral preferences, we applied the same methodology to detect the initiating foot at the start of each epoch with a lower threshold of 1.5 m/s^2^, providing a baseline for normal locomotion initiation.

### Turning

Turning was defined as any instance when in-plane body orientation as derived from the waist tracker crossed a line perpendicular to bush-shelter axis. Based on this definition, we calculated turning count, turning time, turning position, and turning direction. To detect this time point, we first derived the body orientation vector in the horizontal plane, computed as *u*(*t*) = [cos θ_*t*_ sin θ_*t*_], where θ is the yaw angle. A median filter with a window size of 15 (i.e. 300 ms) was applied to remove spike outliers due to sudden jump of trackers data. We then identified the zero-crossings of the first component of the body orientation vector, *u_1_*, which represent potential turning events. To attenuate misdetections due to brief sudden jump in tracker data, we removed zero-crossings that were less than 5 samples (i.e. 100 ms) apart, resulting in the set of filtered zero-crossings. The turning direction was determined by examining the signs of the vector components *u_1_* and *u_2_*, and was assigned as ‘L’ for left turns and ‘R’ for right turns.

### Speed and acceleration profile

Speed and acceleration were computed as the L2 norms of the first and second derivatives of the waist position in three-dimensional space, respectively. Each derivative was smoothed using the same low-pass filter during pre-processing to minimize numerical noise.

### Distance covered

Distance covered represents the total path length traversed by the participant during a specified period, primarily before and after threat appearance. This metric was calculated as the cumulative Euclidean distance between consecutive position measurements of waist tracker in the horizontal plane. Let *p_i_* = (*x_i_,y_i_*) represent the position of the person at time step *i*, where *x_i_* and *y_i_* correspond to the position in x-axis and position in z-axis in the raw file, respectively. The total distance *D* is then computed as the sum of the distances between each pair of consecutive points as depicted below:

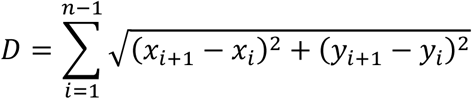

where *n* is the total number of position measurements. This method accounts for the subject’s movement path, including any deviations or non-linear trajectories, providing a more accurate representation of the total distance covered compared to simple start-to-end displacement calculations.

### Foraging posture

Foraging posture was defined as Euclidian distance between two feet during foraging phase. This distance is directly related to wide stance, and indirectly to lower centre of mass.

### Misdirected flight

Misdirected flights are trajectories to a non-shelter location. This was defined by the end position of participant outside the 2 m radius from the shelter and deviates in more than 1 SD from max deviation of trajectory to shelter (escaped trajectories).

### Statistical analysis

The large number of potential hypotheses posed a substantial multiple comparison problem. We opted for a rigorous exploration-confirmation approach. Data exploration was performed in experiment E1. Only significant results within E1 (without correction for multiple comparison) were taken to the confirmation step, and were replicated in independent experiment E2 with Holm-Bonferroni correction for multiple comparison across the entire set of hypotheses. Supporting and exploratory analyses for which only data from one experiment was available are marked as such. The types of tests used are stated in supplementary tables S2-6.

---

**Fig. S1.**
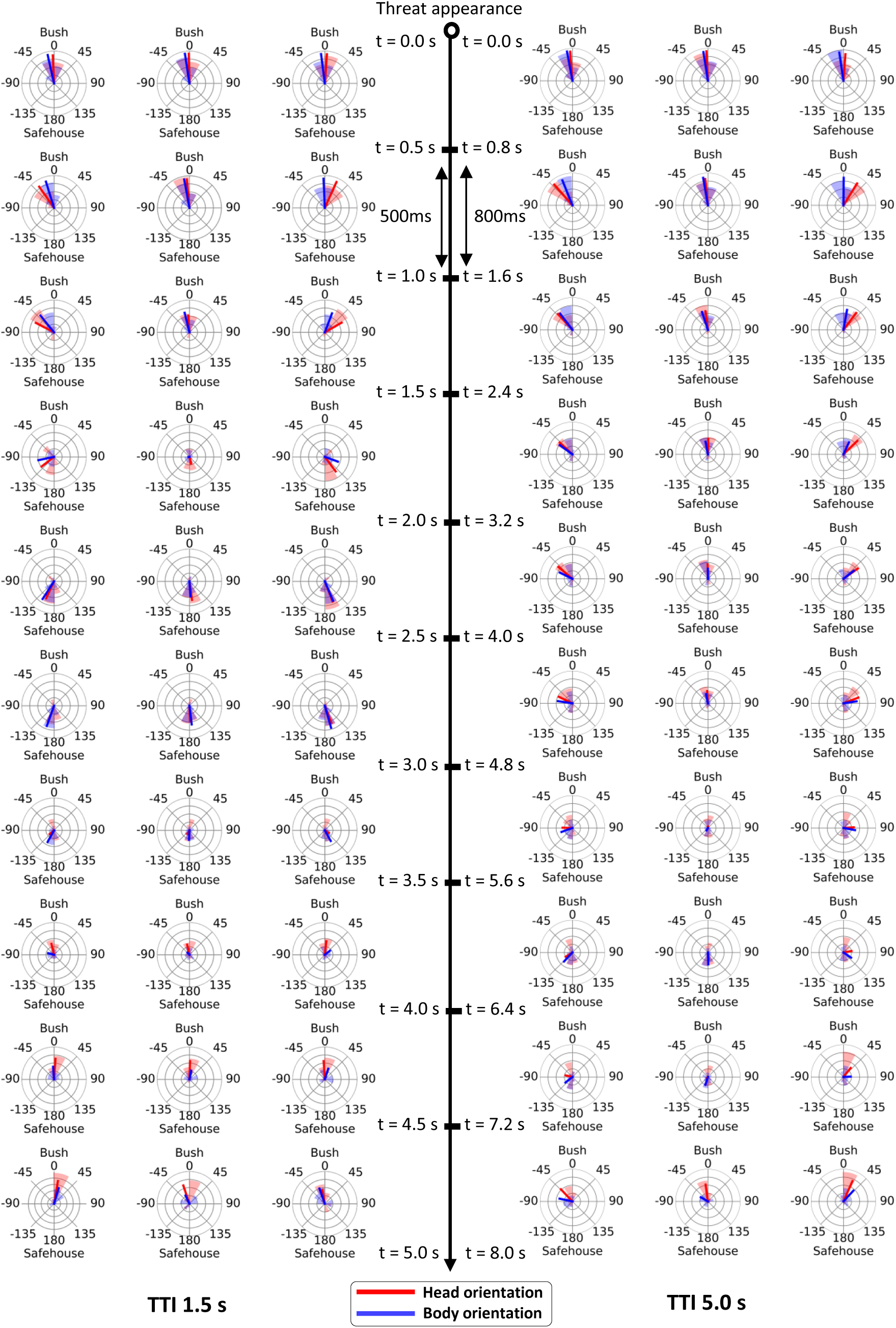
Circular histogram of head and body direction. Relative circular histogram of head direction (red shade) and body direction (blue shade) per 500 ms/800 ms (TTI 1.5 s/TTI 5.0 s) of time after threat appearance relative to threat attack angle (left to right: −45°, 0°, 45°). Red lines show the circular mean value of head direction. Blue lines show the circular mean value of body direction. Bin size is set to 30° and each concentric circle represent a 15% proportion/0.25 vector length for the mean direction.

**Fig. S2.**
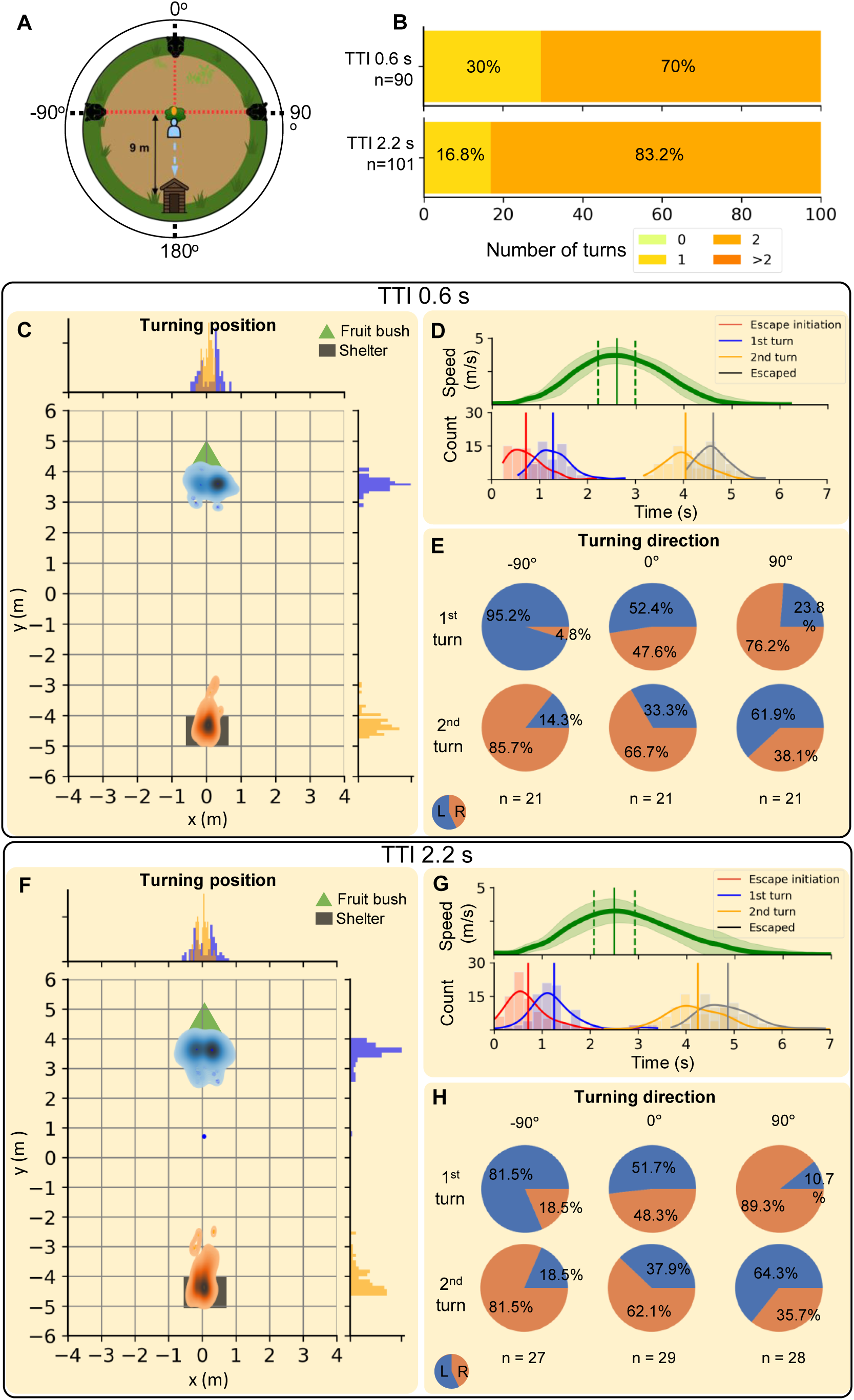
Temporal and spatial characteristics of escape to shelter in E3. (A) Schematic view of the scenario setup. Attack angle is from left (−90°), front (0°) or right (90°). (B) Number of body turns after threat appearance. The dominant movement sequence entails two body turns. (C, F) Distribution of the turning position (position of the waist tracker located on the back of the participant at the time point of turning beyond the x-axis), pooled across all attack angles. Blue points/density shows the first turn and orange the second turn. (D, G) Speed profile and temporal sequence of events following threat appearance at t = 0 s, pooled across all attack angles. The speed profile shows mean (thick green line) and ± 1 SD (shaded area), as well as mean timing of peak speed (vertical line) and its standard deviation (vertical dashed line). For the event sequence, thick vertical lines represent means, and the distribution represents a kernel density estimate. Color-coded events are: escape latency (red), first body turn (blue), second body turn (yellow), and successful entry into the safehouse (black). (E, H) Proportion of turning direction in relation to threat attack angles for the first turn (first row pie chart) and the second turn (second row pie chart). L indicates left body turn, R indicates right body turn, and n indicates number of trials for each condition. Panel C, D, E for TTI 0.6 s, and F, G, H for TTI 2.2 s.

**Fig. S3.**
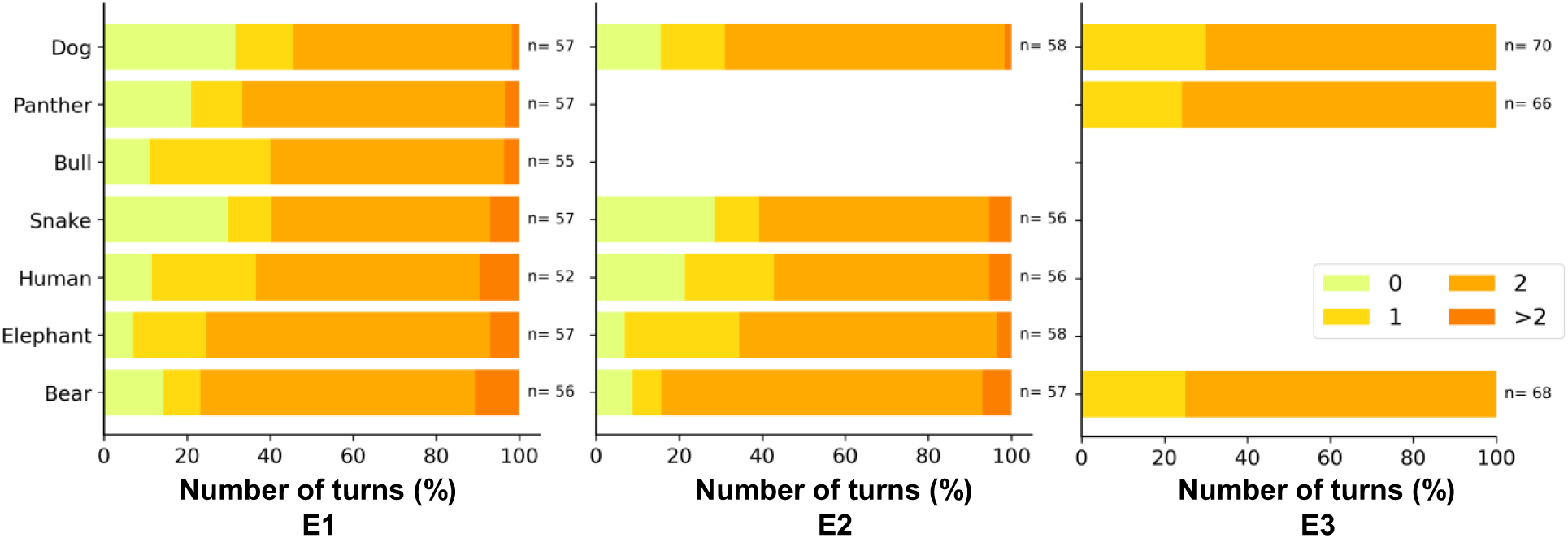
Movement sequence for each threat as indicated by the number of turns as a proportion of all attack epochs.

**Fig. S4.**
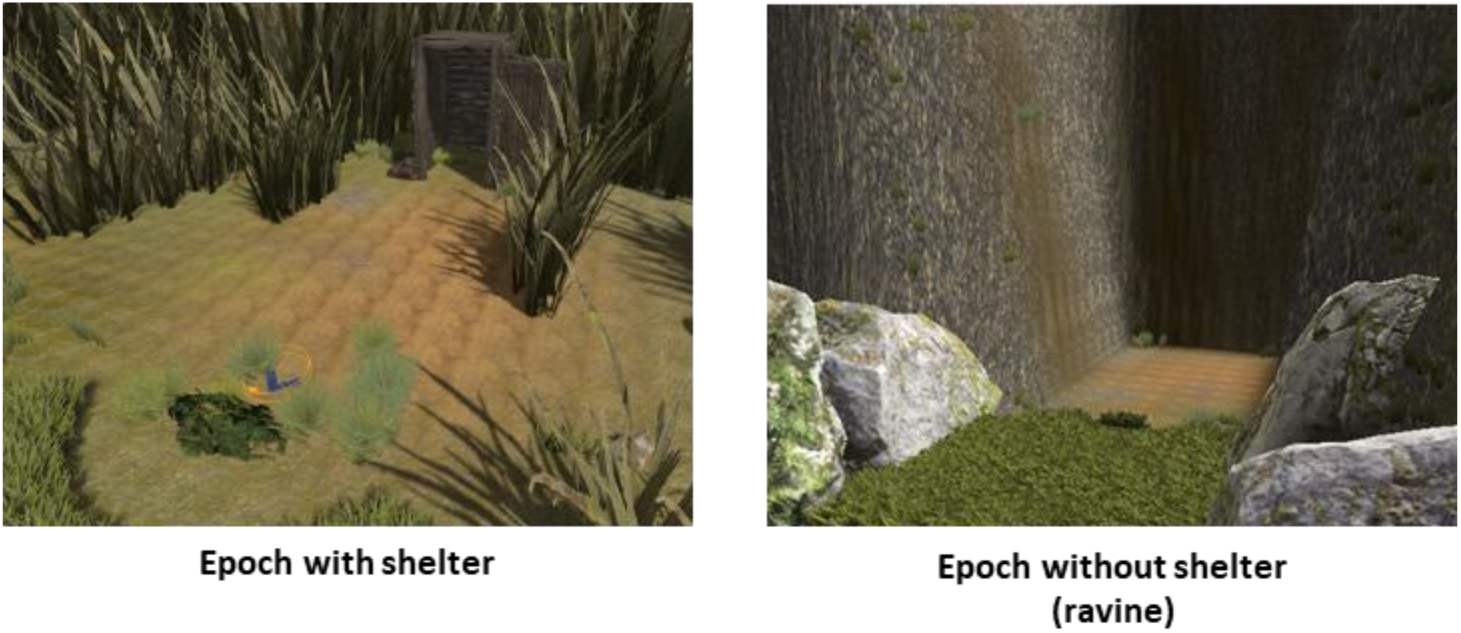
Rendered scenario with and without shelter.

**Table S1.**
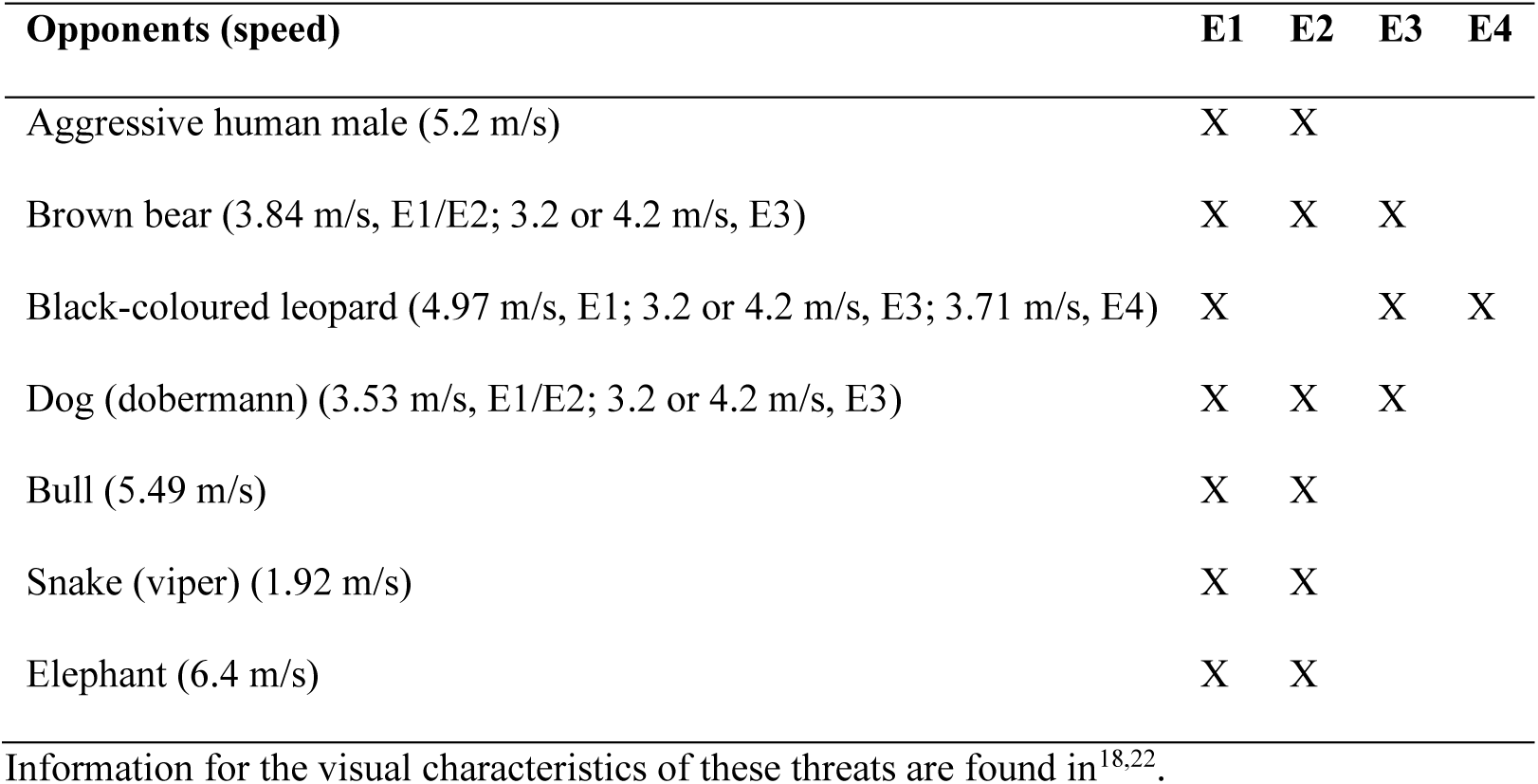
Opponents used in experiments 1-4. Speeds refer to the average speed during the entire attack trajectory to the foraging position, including initial acceleration and deceleration during the attack animation.

**Table S2.** Statistical results of exploration-confirmation approach E1 & E2 and further validation in E3. Long TTI are 5.0 s/2.2 s for E1&E2/E3, and short TTI are 1.5 s/0.6 s for E1&E2/E3. Attack angles are ±45° for E1 and E2, ±90° for E3. N.A. = Not available. Unless otherwise stated, parameter statistics are derived from linear mixed-effects models. All results from linear-mixed effects models were replicated in a robustness analysis with simple t-tests across all participants. E1 was used for data exploration and hypothesis generation; p-values are included for illustration only. All hypothesis tests in E2 and E3 were significant after Holm-Bonferroni^58^ correction for the number of hypotheses tested.

| Dependent variable | Contrast | E1<br>(test statistic, p-value) | E2<br>(test statistic, p-value) | E3<br>(test statistic, p-value) |
| --- | --- | --- | --- | --- |
| Body turning direction (towards/away from threat) | $\pm 45^\circ/\pm 90^\circ$ attack angle, binomial test against 50% ($\hat{p}$: proportion of turns towards threat) | $\hat{p}=0.91$, $n=181$, $p < .0001$ | $\hat{p}=0.77$, $n=128$, $p < .0001$ | $\hat{p}=0.84$, $n=106$, $p < .0001$ |
| Lateralization bias (towards turns in left vs. right attack) | $+45^\circ$ vs $-45^\circ/+90^\circ$ vs $-90^\circ$ attack angle, towards vs. away turns, chi-square test | Cramér's $V=0.82$, $\chi^2=120.28$, $df=1$, $p < .0001$ | $V=0.55$, $\chi^2=38.52$, $df=1$, $p < .0001$ | $V=0.68$, $\chi^2=58.99$, $df=1$, $p < .0001$ |
| Escape strategy | Threat type, bear & elephant vs. dog, snake, human, chi-square test | $\chi^2=6.05$, $df=1$, $p=.0139$ | $\chi^2=8.47$, $df=1$, $p=.0036$ | N.A. (different set of threats) |
| Mean speed | Short TTI, successful vs. unsuccessful escape | $t(182.57)=12.63$, $p < .0001$ | $t(110.23)=11.17$, $p < .0001$ | N.A. (low occurrence of unsuccessful escape) |
| | Long TTI, successful vs. unsuccessful escape | $t(189.99)=9.82$, $p < .0001$ | $t(126.62)=9.82$, $p < .0001$ | N.A. (no unsuccessful escape) |
| Peak speed | Short TTI, successful vs. unsuccessful escape | $t(190.97)=10.82$, $p < .0001$ | $t(137.50)=8.58$, $p < .0001$ | N.A. (low occurrence of unsuccessful escape) |
| | Long TTI, successful vs. unsuccessful escape | $t(192.47)=4.56$, $p < .0001$ | $t(126.77)=3.80$, $p=.0002$ | N.A. (no unsuccessful escape) |
| Time to peak speed | Long TTI, successful vs. unsuccessful escape | $t(190.21)=-3.53$, $p=.0005$ | $t(132.82)=-4.91$, $p < .0001$ | N.A. (no unsuccessful escape) |
| Mean acceleration magnitude | Short TTI, successful vs. unsuccessful escape | $t(189.49)=8.67, p < .0001$ | $t(134.55)=5.31, p < .0001$ | N.A. (low occurrence of unsuccessful escape) |
| | Long TTI, successful vs. unsuccessful escape | $t(189.53)=6.39, p < .0001$ | $t(125.51)=7.04, p < .0001$ | N.A. (no unsuccessful escape) |
| Peak acceleration magnitude | Short TTI, successful vs. unsuccessful escape | $t(190.76)=3.33, p = .0010$ | $t(135.72)=2.61, p = .0101$ | N.A. (low occurrence of unsuccessful escape) |
| Time to peak acceleration | Long TTI, successful vs. unsuccessful escape | $t(195.87)=-3.26, p = .0013$ | $t(134.48)=-4.88, p < .0001$ | N.A. (no unsuccessful escape) |
| Escape initiation time (escape latency) | Short TTI, successful vs. unsuccessful escape | $t(185.00)=-5.01, p < .0001$ | $t(135.99)=-3.74, p = .0002$ | N.A. (low data count, unsuccessful escape) |
| | Long TTI, successful vs. unsuccessful escape | $t(177.64)=-3.02, p = .0029$ | $t(117.41)=-4.67, p < .0001$ | N.A. (no unsuccessful escape) |
| Backward escape leads to getting caught | Short TTI, first turn time in successful escape vs. caught time in unsuccessful escape | $t(174.61)=-36.48, p < .0001$ | $t(122.30)=-39.68, p < .0001$ | N.A. (no backward escape) |
| | Long TTI, first turn time in successful escape vs. caught time in unsuccessful escape | $t(165.55)=-21.03, p < .0001$ | $t(117.56)=-22.70, p < .0001$ | N.A. (no backward escape) |
| Distance covered | Before threat appearance, shelter vs. no shelter | N.A. | $t(312.68)=-10.68, p < .0001$ | N.A. |

**Table S3.** Determinants of unsuccessful escape. Results are presented as mean **±** SD for E1 and E2 combined.

| Time to impact (TTI) | Determinants | Unsuccessful escape | Successful escape |
| --- | --- | --- | --- |
| 1.5 s | Mean speed (m/s) | 0.80 ± 0.35 | 1.27 ± 0.13 |
| | Peak speed (m/s) | 1.85 ± 0.71 | 2.81 ± 0.39 |
| | Mean acceleration magnitude (m/s<sup>2</sup>) | 1.47 ± 0.56 | 2.05 ± 0.35 |
| | Escape latency (s) | 1.06 ± 0.62 | 0.72 ± 0.43 |
| 5.0 s | Mean speed (m/s) | 0.47 ± 0.29 | 0.85 ± 0.17 |
| | Peak speed (m/s) | 1.71 ± 0.98 | 2.29 ± 0.58 |
| | Time to peak speed (s) | 5.76 ± 1.77 | 4.61 ± 1.54 |
| | Mean acceleration magnitude (m/s<sup>2</sup>) | 0.96 ± 0.5 | 1.40 ± 0.30 |
| | Time to peak acceleration (s) | 5.45 ± 2.02 | 3.99 ± 2.08 |
| | Escape latency (s) | 3.32 ± 1.75 | 2.10 ± 1.51 |

**Table S4.**
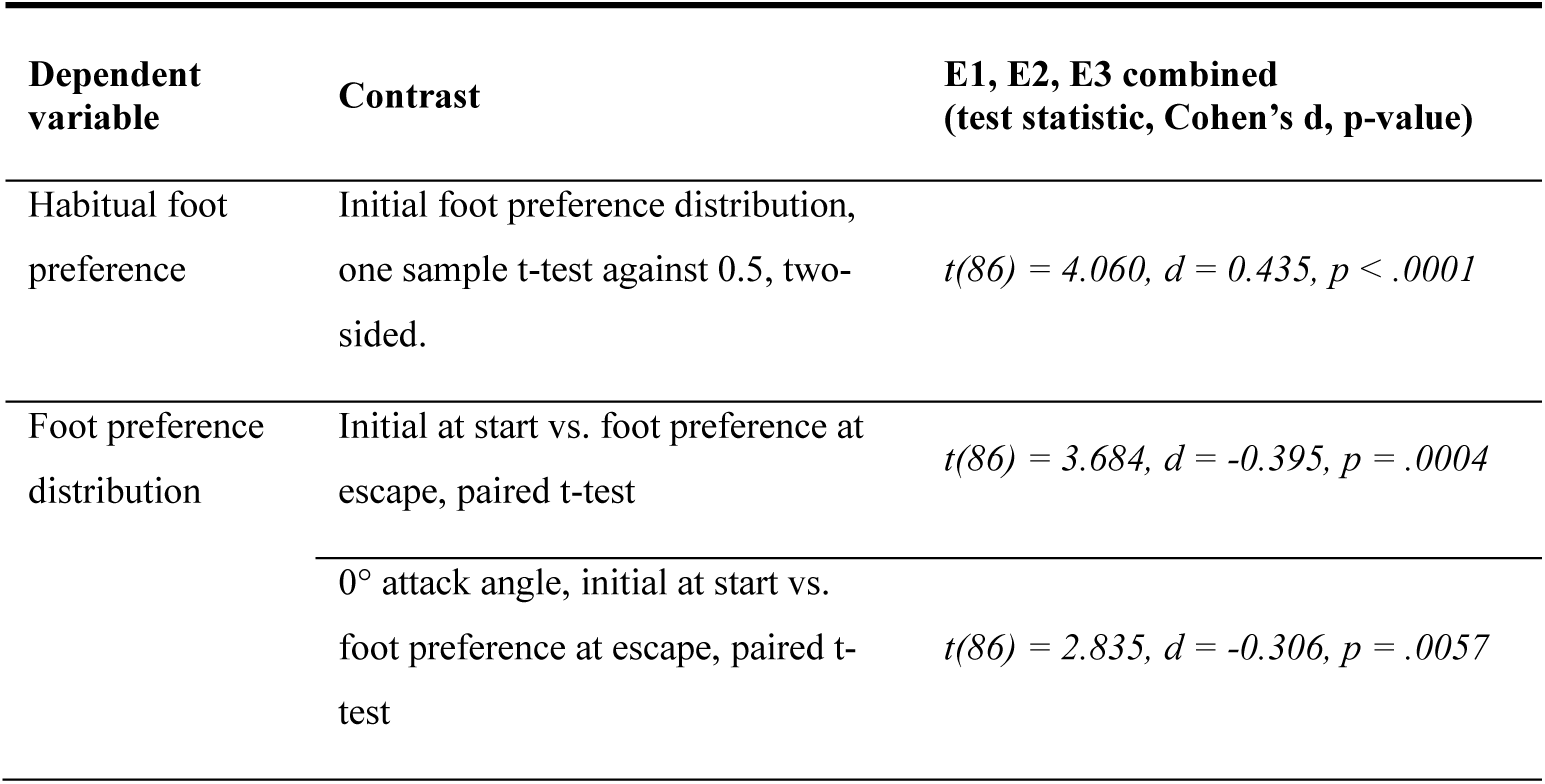
Analysis of foot preference. For foot preferences, due to incomplete or missing foot tracker data, we pooled all participants (N=87) for the analysis of foot preference. Before doing so, we confirmed that there was no detectable batch effect/experiment to experiment variation. This justified treating the combined dataset as a single cohort for our primary inferential test for foot preferences.

**Table S5.** Determinants of escape foot preference. Results from a mixed-effect logistic regression model with the formula: Escape foot preference ∼ Turning direction + Threat attack angle + Habitual foot preference + (1 | participant) + (1 | experiment). The estimated variance of participant intercepts was .84 (SD=.92), suggesting that some participants were consistently more likely to choose the right foot. Non-zero intercept here is a baseline preference for the left foot. Data from E1, E2, and E3 (N=86, n=465).

| Fixed Effects | Coefficient (S.E.) | z-value | p-value |
| --- | --- | --- | --- |
| Intercept | -0.5433 (0.245) | -2.21 | .0269* |
| Turning direction (Right) | 0.9038 (0.317) | 2.85 | .0044** |
| Attack angle (Right) | -0.1164 (0.307) | -0.38 | .7043 |
| Initial foot preference | 0.1727 (0.242) | 0.71 | .4763 |

\*p<.05, \*\*p<.01

**Table S6.** Analysis of foraging posture in E4. Results from linear mixed-effects models with the formula: DV ∼ Contrast + (1 | participant).

| Dependent variable (DV) | Contrast | $\beta \pm SE, t(df), p$ |
| --- | --- | --- |
| Foraging posture<br>(Euclidian feet distance) | E4 vs. E1 | $0.100 \pm 0.034, t(101)=2.99, p = .0096$ |
| | E4 vs. E2 | $0.167 \pm 0.037, t(104)=4.55, p < .0001$ |
| | E1 vs. E2 | $0.066 \pm 0.041, t(103)=1.62, p = .241$ |
| Foraging posture<br>(Euclidian feet distance) | Escaped vs. Caught<br>(2, 3, 4 m threat distance) | $0.091 \pm 0.034, t(275.28)=2.68, p = .0077$ |

## Supplementary Video

This video includes two camera views, a VR view, a 2D trajectory plot, a 3D stick-figure animation, and an event sequence plot, illustrating the distinct motor sequence of human escape to shelter.

## Notes

https://osf.io/2b3k7/

https://osf.io/pmdjt/

https://xip.uclb.com/product/vrthreat-toolkit-for-unity

### Competing Interest Statement

The authors have declared no competing interest.

### Summary of Updates

Results updated. Discussions are updated. Supplemental files updated.

## References

1. Domenici, P., Blagburn, J. M. & Bacon, J. P. Animal escapology I: Theoretical issues and emerging trends in escape trajectories. J. Exp. Biol. 214, 2463–2473 (2011).

2. Ledoux, J. & Daw, N. D. Surviving threats: Neural circuit and computational implications of a new taxonomy of defensive behaviour. Nat. Rev. Neurosci. 19, 269–282 (2018).

3. Cooper, W. E. & Blumstein, D. T. Escaping From Predators. Escaping from Predators: An Integrative View of Escape Decisions (Cambridge University Press, 2015). doi:10.1017/CBO9781107447189.

4. Evans, D. A., Stempel, A. V., Vale, R. & Branco, T. Cognitive Control of Escape Behaviour. Trends Cogn. Sci. 23, 334–348 (2019).

5. Domenici, P., Blagburn, J. M. & Bacon, J. P. Animal escapology II: Escape trajectory case studies. J. Exp. Biol. 214, 2474–2494 (2011).

6. Wilson, R. S., Husak, J. F., Halsey, L. G. & Clemente, C. J. Predicting the Movement Speeds of Animals in Natural Environments. Integr. Comp. Biol. 55, 1125–1141 (2015).

7. Wilson, A. M. et al. Biomechanics of predator-prey arms race in lion, zebra, cheetah and impala. Nature 554, 183–188 (2018).

8. Stankowich, T. & Coss, R. G. Effects of risk assessment, predator behavior, and habitat on escape behavior in Columbian black-tailed deer. Behav. Ecol. 18, 358–367 (2007).

9. Walker, J. A., Ghalambor, C. K., Griset, O. L., McKenney, D. & Reznick, D. N. Do faster starts increase the probability of evading predators? Funct. Ecol. 19, 808–815 (2005).

10. Kimura, H. et al. Escaping from multiple visual threats: modulation of escape responses in Pacific staghorn sculpin (Leptocottus armatus). J. Exp. Biol. 225, (2022).

11. Branco, T. & Redgrave, P. The Neural Basis of Escape Behavior in Vertebrates. Annu. Rev. Neurosci. 43, 417–439 (2020).

12. Domenici, P. & Hale, M. E. Escape responses of fish: A review of the diversity in motor control, kinematics and behaviour. J. Exp. Biol. 222, (2019).

13. Kawabata, Y. et al. Multiple preferred escape trajectories are explained by a geometric model incorporating prey’s turn and predator attack endpoint. Elife 12, 1–29 (2023).

14. Moore, T. Y., Cooper, K. L., Biewener, A. A. & Vasudevan, R. Unpredictability of escape trajectory explains predator evasion ability and microhabitat preference of desert rodents. Nat. Commun. 8, 440 (2017).

15. Bach, D. R. Cross-species anxiety tests in psychiatry: pitfalls and promises. Mol. Psychiatry 27, 154–163 (2022).

16. Bach, D. R., Moutoussis, M., Bowler, A. & Dolan, R. J. Predictors of risky foraging behaviour in healthy young people. *Nat*. Hum. Behav. 4, 832–843 (2020).

17. Slater, M., Spanlang, B., Sanchez-Vives, M. V. & Blanke, O. First person experience of body transfer in virtual reality. PLoS One 5, 1–9 (2010).

18. Brookes, J., Hall, S., Frühholz, S. & Bach, D. R. Immersive VR for investigating threat avoidance: The VRthreat toolkit for Unity. Behav. Res. Methods 56, 5040–5054 (2023).

19. Baker, C., Pawling, R. & Fairclough, S. Assessment of threat and negativity bias in virtual reality. Sci. Rep. 10, 1–10 (2020).

20. Binder, F. P. & Spoormaker, V. I. Quantifying Human Avoidance Behavior in Immersive Virtual Reality. Front. Behav. Neurosci. 14, 1–17 (2020).

21. Topalovic, U. et al. Wireless Programmable Recording and Stimulation of Deep Brain Activity in Freely Moving Humans. Neuron 108, 322–334.e9 (2020).

22. Sporrer, J. K. et al. Functional sophistication in human escape. iScience 26, 108240 (2023).

23. Brookes, J., Warburton, M., Alghadier, M., Mon-Williams, M. & Mushtaq, F. Studying human behavior with virtual reality: The Unity Experiment Framework. Behav. Res. Methods 52, 455–463 (2020).

24. Driver, P. M. & Humphries, D. A. Protean displays as inducers of conflict. Nature 226, 968–969 (1970).

25. Edut, S. & Eilam, D. Protean behavior under barn-owl attack: Voles alternate between freezing and fleeing and spiny mice flee in alternating patterns. Behav. Brain Res. 155, 207–216 (2004).

26. Richardson, G., Dickinson, P., Burman, O. H. P. & Pike, T. W. Unpredictable movement as an anti-predator strategy. Proc. R. Soc. B Biol. Sci. 285, (2018).

27. Cantalupo, C., Bisazza, A. & Vallortigara, G. Lateralization of predator-evasion response in a teleost fish (Girardinus falcatus). Neuropsychologia 33, 1637–1646 (1995).

28. Kurvers, R. H. J. M. et al. The Evolution of Lateralization in Group Hunting Sailfish. Curr. Biol. 27, 521–526 (2017).

29. Fisher, R. A. Statistical Methods for Research Workers. (Oliver and Boyd, 1925).

30. Ruxton, G. D. & Neuhäuser, M. Good practice in testing for an association in contingency tables. Behav. Ecol. Sociobiol. 64, 1505–1513 (2010).

31. Shamash, P. et al. Mice learn multi-step routes by memorizing subgoal locations. Nat. Neurosci. 24, 1270–1279 (2021).

32. Tomsic, D., Sztarker, J., De Astrada, M. B., Oliva, D. & Lanza, E. The predator and prey behaviors of crabs: From ecology to neural adaptations. J. Exp. Biol. 220, 2318–2327 (2017).

33. Ellard, C. G. & Goodale, M. A. A functional analysis of the collicular output pathways: a dissociation of deficits following lesions of the dorsal tegmental decussation and the ipsilateral collicular efferent bundle in the Mongolian gerbil. Exp. Brain Res. 71, 168–172 (1988).

34. Vale, R., Evans, D. A. & Branco, T. Rapid Spatial Learning Controls Instinctive Defensive Behavior in Mice. Curr. Biol. 27, 1342–1349 (2017).

35. Campagner, D. et al. A cortico-collicular circuit for orienting to shelter during escape. Nature 613, 111–119 (2023).

36. Lippolis, G., Bisazza, A., Rogers, L. J. & Vallortigara, G. Lateralisation of predator avoidance responses in three species of toads. *Laterality Asymmetries Body*, Brain Cogn. 7, 163–183 (2002).

37. Phillips, C. J. C., Oevermans, H., Syrett, K. L., Jespersen, A. Y. & Pearce, G. P. Lateralization of behavior in dairy cows in response to conspecifics and novel persons. J. Dairy Sci. 98, 2389–2400 (2015).

38. Rogers, L. J. Evolution of Hemispheric Specialization: Advantages and Disadvantages. Brain Lang. 73, 236–253 (2000).

39. Vallortigara, G. The evolutionary psychology of left and right: Costs and benefits of lateralization. Dev. Psychobiol. 48, 418–427 (2006).

40. Henry, S. M., Fung, J. & Horak, F. B. Effect of stance width on multidirectional postural responses. J. Neurophysiol. 85, 559–570 (2001).

41. Goodworth, A. D. & Peterka, R. J. Influence of stance width on frontal plane postural dynamics and coordination in human balance control. J. Neurophysiol. 104, 1103–1118 (2010).

42. Otsuka, M., Kurihara, T. & Isaka, T. Effect of a wide stance on block start performance in sprint running. PLoS One 10, (2015).

43. Brochard, J., Dayan, P. & Bach, D. R. Critical intelligence: Computing defensive behaviour. Neurosci. Biobehav. Rev. 174, 106213 (2025).

44. Woods, D. L., Wyma, J. M., Yund, E. W., Herron, T. J. & Reed, B. Factors influencing the latency of simple reaction time. Front. Hum. Neurosci. 9, 1–12 (2015).

45. Haith, A. M., Pakpoor, J. & Krakauer, J. W. Independence of movement preparation and movement initiation. J. Neurosci. 36, 3007–3015 (2016).

46. Craske, M. G. et al. Anxiety disorders. Nat. Rev. Dis. Prim. 3, 17024 (2017).

47. Bach, D. R. & Dayan, P. Algorithms for survival: A comparative perspective on emotions. Nat. Rev. Neurosci. 18, 311–319 (2017).

48. van Ast, V. A., Klumpers, F., Grasman, R. P. P. P., Krypotos, A. M. & Roelofs, K. Postural freezing relates to startle potentiation in a human fear-conditioning paradigm. Psychophysiology 59, 1–20 (2022).

49. Yilmaz, M. & Meister, M. Report Rapid Innate Defensive Responses of Mice to Looming Visual Stimuli. Curr. Biol. 23, 2011–2015 (2015).

50. Shang, C. et al. Divergent midbrain circuits orchestrate escape and freezing responses to looming stimuli in mice. Nat. Commun. 9, 1–17 (2018).

51. Lenzi, S. C. et al. Threat history controls flexible escape behavior in mice. Curr. Biol. 32, 2972–2979.e3 (2022).

52. Himmels, C. et al. Validating risk behavior in driving simulation using naturalistic driving data. Transp. Res. Part F Traffic Psychol. Behav. 107, 710–725 (2024).

53. Horsak, B. et al. Overground Walking in a Fully Immersive Virtual Reality: A Comprehensive Study on the Effects on Full-Body Walking Biomechanics. Front. Bioeng. Biotechnol. 9, 1–10 (2021).

54. Krakauer, J. W., Ghazanfar, A. A., Gomez-Marin, A., MacIver, M. A. & Poeppel, D. Neuroscience Needs Behavior: Correcting a Reductionist Bias. Neuron 93, 480–490 (2017).

55. Niehorster, D. C., Li, L. & Lappe, M. The accuracy and precision of position and orientation tracking in the HTC vive virtual reality system for scientific research. Iperception. 8, 1–23 (2017).

56. Caserman, P., Garcia-Agundez, A., Konrad, R., Göbel, S. & Steinmetz, R. Real-time body tracking in virtual reality using a Vive tracker. Virtual Real. 23, 155–168 (2019).

57. Kavanagh, J. J. & Menz, H. B. Accelerometry: A technique for quantifying movement patterns during walking. Gait Posture 28, 1–15 (2008).

58. Holm, S. A Simple Sequentially Rejective Multiple Test. Scand. J. Stat. 6, 65–70 (1979).

